# Phenotypic and physiological responses to salt exposure in *Sorghum* reveal diversity among domesticated landraces

**DOI:** 10.1101/848028

**Authors:** Ashley N. Henderson, Philip M. Crim, Jonathan R. Cumming, Jennifer S. Hawkins

## Abstract

Soil salinity negatively impacts plant function, development, and yield. *Sorghum bicolor* is a staple crop known to be drought tolerant, to have adapted to a variety of conditions, and to contain significant standing genetic diversity, making it an exemplary species to study phenotypic and physiological variation in salinity tolerance. In our study, a diverse group of sorghum landraces and accessions was first rank-ordered for salinity tolerance and then individuals spanning a wide range of response were analyzed for foliar proline and ion accumulation. We found that, while proline is often a good indicator of osmotic adjustment and is historically associated with increased salt tolerance, proline accumulation in sorghum reflects stress-response injury rather than acclimation. When combining ion profiles with growth responses and stress tolerance indices, the variation observed in tolerance was similarly not a sole result of Na^+^ accumulation, but rather reflected accession-specific mechanisms that may integrate these and other metabolic responses. When we compared variation in tolerance to phylogenetic relationships, we conclude that the most parsimonious explanation for the variation observed among accessions is that salinity tolerance was acquired early during domestication and was subsequently maintained or lost in diverged lineages during improvement in areas that vary in soil salinity.

## INTRODUCTION

Soil salinity is a major constraint to agricultural crop productivity, limiting the provision of food, fuel, and fiber to large portions of the world’s population. Soil salinity, defined as concentrations of soluble salts above 40 mM sodium chloride (NaCl) or greater than 4 dSm^-1^ electrical conductivity (Jamil *et al*., 2011; Shrivastava & Kumar, 2015), is a global problem affecting more than 20% of the irrigated land used for agriculture (Qadir *et al*., 2014). Salts increase in soils naturally through the rise and ingression of sea water (Abrol *et al*., 1988; Singh, 2015; Liu *et al*., 2017), weathering of soil parent material (Abrol *et al*., 1988), and low precipitation accompanied by high surface evaporation (Chhabra, 1996; Shrivastava & Kumar, 2015; Singh, 2015). Anthropogenic factors, such as irrigation with saline water, inadequate field drainage, and over application of animal waste, also result in increased soluble salts in agricultural soils (Munns & Tester, 2008; Thomson *et al*., 2010; Singh, 2015; Lemanowicz & Bartkowiak, 2017).

Increased salinity negatively impacts plant function and development through both osmotic and ionic effects (Negrão *et al*., 2017). In the osmotic phase, salinity impedes plant water acquisition. Water uptake is disrupted even when soils contain adequate moisture due to the lower soil water potentials compared to plant osmotic potentials. This imbalance inhibits water extraction by plant roots, simulating drought-like conditions (Munns & Tester, 2008; Negrão *et al*., 2017). In order to alleviate the osmotic effects associated with salinity, plants produce compatible solutes, such as amino acids, amines, betaines, organic acids, sugars, and poylols, that aid in osmotic adjustment and assist in the movement of water into the plant (Parihar *et al*., 2015). In the ion-dependent phase, ions such as Na^+^ and Cl^-^ enter the plant, accumulate to toxic levels in cells, and disrupt normal metabolic function (Munns & Tester, 2008). Plant ion transport systems function to exclude toxic ions from the cytoplasm, through either extrusion or compartmentalization, in order to maintain homeostasis (Munns & Tester, 2008).

Various plant responses result from both ion-independent and dependent phases. Key growth responses to osmotic stress include decreased leaf and root growth due to lack of turgor (Munns, 2005). Leaf growth is affected to a greater extent than root growth, resulting in a decreased shoot to root ratio (Negrão *et al*., 2017). This is an adaptive response because, with decreased leaf biomass, less water is lost from the plant canopy resulting in less uptake from the soil (Iqbal *et al*., 2014), ultimately reducing salt concentrations at the root surface (Munns, 2010). Toxic ion buildup in leaves affects ion homeostasis and photosynthesis, resulting in premature leaf senescence (Munns, 1993, 2002). As ions accumulate, Na^+^ specifically disrupts the uptake and distribution of K^+^, an essential ion for basic biological functions such as stomatal opening and enzyme activity (Tari *et al*., 2013) or cellular metabolism (Zhu, 2003); however, because salts may be compartmentalized into vacuoles and older leaves, plants can survive the ionic component of salt stress if the rate of new leaf emergence exceeds the rate of leaf death. This enables the plant to continue photosynthesizing and fixing carbon to sustain growth and development (Munns, 2005, 2010). The ability to maintain a high K^+^/Na^+^ ratio is often a strong indication of salt tolerant genotypes (Thomson *et al*., 2010; Mahi *et al*., 2019).

*Sorghum bicolor* (L.) Moench is an African grass that is cultivated for food, fuel, and fiber. Worldwide, it ranks fifth as a contributor to grain production and second as a biofuels feedstock (Wiersema & Dahlberg, 2007). Sorghum thrives in areas that are often not suitable for other crops and requires minimal human input while delivering high yields (Mullet *et al*., 2014). Given these traits, sorghum provides a model system for studying the complex basis of salt tolerance because it is relatively drought tolerant (Mullet *et al*., 2014; Fracasso *et al*., 2016; McCormick *et al*., 2018) and, as with drought stress, salinity stress results in osmotic imbalance (Munns & Tester, 2008). Additionally, previous studies have shown significant genetic diversity within domesticated sorghum (landraces and improved varieties), making it an ideal system to discern the standing variation associated salinity response.

Here, we evaluated the variation in whole-plant response to salt exposure in a diverse panel of sorghum accessions and wild relatives. Specifically, we include a hybrid species, three wild progenitors, and a variety of cultivated landraces to evaluate the association between genotypic diversity with salinity tolerance. Our findings indicate that landrace is not the primary determinant of salinity tolerance. We observed racial structure influencing growth traits, but a lack of association between landrace and key physiological responses to NaCl. Therefore, we further compared our tolerance groupings with the known phylogenetic relationships outlined by Mace et al. (2013). Together, our results suggest that salinity tolerance originated early during domestication and was maintained and/or lost throughout improvement in areas that vary in soil salinity.

## MATERIALS AND METHODS

### Plant Material

There are five landraces of sorghum (bicolor, kafir, guinea, caudatum, and durra) that are classified based on morphology (Shehzad *et al*., 2009) and reflect different geographical regions of adaptation (Price *et al*., 2005; Morris *et al*., 2013; Mace *et al*., 2013; Mullet *et al*., 2014; Smith *et al*., 2019). There are also 10 intermediate landraces that are a combination of the five landraces (Oliveira *et al*., 1996; Price *et al*., 2005). Sorghum was improved in a diversity of environments and is a staple grain in various regions (Smith & Frederiksen, 2000). Because *Sorghum bicolor* was originally domesticated c. 5,000 years ago in eastern Africa (Wendorf *et al*., 1992; Mace *et al*., 2013; Winchell *et al*., 2017; Smith *et al*., 2019), we hypothesized that varying degrees of sensitivity and tolerance to NaCl may exist in the different landraces. In this study, we included 21 diverse *Sorghum* accessions representative of the different landraces (**Table 1**). In addition, the accessions included in this study display important agricultural traits and are lines included in the Sorghum Association Mapping population (Jordan *et al*., 2011). These serve as valuable resources when dissecting complex traits, such as salinity tolerance.

**Table 1.** Summary of sorghum accessions. Sorghum accessions and associated information (identification code used to reference accessions throughout the study and landrace). Accession information and landrace information was supplied by GRIN.

| Accession | ID | Landrace | Sorghum Association Panel |
| --- | --- | --- | --- |
| PI33027204SD | D-1 | drummondii |  |
| subs. propinquum | P-1 | subs. *propinquum* |  |
| PI57112801SD | Sb-1 | caudatum |  |
| PI53412801SD | Sb-2 | durra | SAP-208 |
| PI52569503SD | Sb-3 | guinea-margaritifera |  |
| PI57613001SD | Sb-4 | kafir | SAP-65 |
| PI53391004SD | Sb-5 | caudatum | SAP-268 |
| PI53383401SD | Sb-6 | caudatum |  |
| PI53379202SD | Sb-7 | caudatum | SAP-140 |
| PI65606902SD | Sb-8 | intermediate (unknown) |  |
| PI58643001SD | Sb-9 | guinea-margaritifera |  |
| PI58574902SD | Sb-10 | durra |  |
| PI56512103SD | Sb-11 | caudatum | SAP-80 |
| PI53413301SD | Sb-12 | durra | SAP-233 |
| PI53375201SD | Sb-13 | caudatum | SAP-127 |
| PI65361702SD | Sb-14 | intermediate (unknown) | SAP-73 |
| PI61353602SD | Sb-15 | durra-caudatum | SAP-74 |
| PI56351602SD | Sb-16 | durra-caudatum |  |
| PI60933601SD | Sb-17 | intermediate (unknown) |  |
| PI65602902SD | Sb-18 | durra | SAP-37 |
| Tx7000 | Tx-1 | durra |  |
| PI22609603SD | V-1 | subs. *verticilliflorum* |  |
| PI30011903SD | V-2 | subs. *verticilliflorum* |  |

Note: Tx-1 and Sb-6 were excluded from the study

All seeds were obtained from the *Germplasm Resources Information Network (GRIN)*. Landrace information was provided by GRIN and arbitrary codes were assigned and used to reference specific accessions throughout this study (**Table 1**).

### NaCl Exposure

A pilot study, in which five randomly selected accessions were exposed to increasing salt concentrations, was used to determine an appropriate experimental treatment level. Replicates were treated with 0 mM, 25 mM, 75 mM, 125 mM, 150 mM, or 200 mM NaCl beginning at the third leaf stage of development and for a period of four weeks. There was a clear reduction in growth and biomass as NaCl increased (**Supplementary Fig. S1**). Because soil is considered to be saline at concentrations greater than 40 mM (Shrivastava & Kumar, 2015) and we observed growth reduction without mortality at 75 mM NaCl, we utilized this concentration for further intensive study.

Twenty seeds of each accession (10 replicates per treatment and a total of two treatments) were germinated in metromix soil in 5 cm × 5 cm × 5 cm planting plugs under 29/24°C day/night temperatures in controlled greenhouse conditions. During germination, all seedlings were misted regularly with non-saline tap water. When 90% of the seedlings were at the third leaf stage of development, seedlings were transplanted into 5 cm × 5 cm × 25 cm treepots (Stuewe and Sons, Tangent, OR, USA) filled with a 1:1 mix of #2 and #4 silica sand. Seedlings were watered with tap water for one-week post-transplant to provide a period of establishment.

After establishment, plants were watered to saturation daily with tap water (control) or tap water containing 75 mM NaCl solution (treatment). Twice each week, all plants were additionally watered to saturation with a 20-10-20 N-P-K fertilizer at a rate of 200 ppm (J.R. Peters, Inc., Allentown, PA, USA). Treatment was carried out for a total of 12 weeks.

### Biomass Measurements

At 12 weeks post treatment, five of the ten replicates were collected for biomass measurements. Biomass samples were cut, bagged, and dried in four different categories (belowground, stem, live leaves (defined as >50% green leaf), dead leaves (defined as <50% green leaf), and tillers]. All biomass samples were dried at 65°C.

Throughout this study, the following terms were used to describe the following tissues: live above ground biomass was the sum of the live stem, live leaves, and live tillers. Dead above ground biomass was the sum of the dead stem, dead leaves, and dead tillers. Total above ground biomass was the sum of live and dead above ground biomass. Percent of live above ground biomass was the ratio of live above ground biomass by the total above ground biomass as a fraction of 100.

### Phenotype Measurements

The remaining five replicates were used for phenotypic measurements. The following phenotypic measurements were recorded after 12 weeks of treatment: total number of leaves, total number of live leaves, percent live leaves (calculated from live leaves and total leaves), mortality (defined as 1 for alive and 0 for dead), and height (cm).

### Physiology Measurements

Physiology measurements were taken at 12 weeks post treatment on the third leaf from the top because it was the oldest living leaf across all plants. The same five replicates used for phenotypic measurements were used for quantification of chlorophyll content (SPAD 502 Plus Chlorophyll Meter, Konica Minolta, Osaka, Japan) and quantification of proline content. Ion profiles were measured on the same five replicates used for biomass measurements. Proline content and ion profiles were quantified on a subset of accessions that showed variation in phenotypic responses. SPAD was recorded on all accessions and replicates.

Foliar sodium and potassium concentrations were determined on microwave-assisted acid digests (MARSXpress, CEM Corporation, Matthews, NC, USA). Leaf tissue was dried for 72 h at 70°C, ground in a CyclotecTM 1093 sample mill (FOSS, Hilleroed, Denmark), and digested in 4 mL of 70% HNO_3_ and 1 mL of 30% H_2_O_2_ (Carrilho *et al*., 2002). Digests were analyzed for elemental concentrations by inductively coupled plasma optical emission spectrometry (ICP-OES) by the Pennsylvania State University Analytical Laboratory (State College, PA, USA). Elemental yields were obtained using ground apple leaves from the National Institute of Standards and Technology and were used to calculate elemental content from the ICP-OES data.

Quantification of proline was determined colorimetrically by comparisons with standards. Following harvest, samples were flash frozen and immediately stored at −80°C. Tissue was ground to a fine powder and 2 mL of 70% ethanol was added to each sample. Samples were incubated at room temperature with continuous agitation for 24 h, after which they were centrifuged and the supernatant was transferred to a new tube. The ground tissue was then re-suspended in fresh 2 mL of 70% ethanol for an additional 24 h at room temperature with agitation. After the second extraction, both 2 mL extracts were combined. Samples were then incubated at 95°C for 20 min with a 1% ninhydrin and 60% acetic acid reaction mix and quantified on a Tecan Infinite® 200 PRO plate reader (Tecan, Grödig, Austria) at 520 nm.

### Statistical Analyses

Salinity tolerance in plants is often defined as the ability of a plant to sustain growth in the presence of salts (Munns, 2010). In our study, several parameters were evaluated and tolerance was defined by the ability to maintain biomass (live and total) when comparing salt exposure to control conditions (Negrão *et al*., 2017).

#### Stress Tolerance (ST)

The stress tolerance value was calculated for SPAD of the oldest living leaf across all plants, percent of live leaves, height (cm), mortality, live aboveground biomass (dry weight in g), dead aboveground biomass (dry weight in g), and root biomass (dry weight in g) as (Negrão *et al*., 2017):

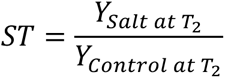

Where Y is a growth-related trait measured at the end of the experiment (T_2_) under control and salt treatments as indicated. The ST value normalizes performance by accession.

#### Relative Decrease in Plant Biomass (RDPB)

The sum of biomass for all tissues separated during a destructive harvest was used to determine the relative decrease in plant biomass (RDPB, Negrão *et al*., 2017) for each accession and landrace. The RDPB describes the reduction of growth in stressed conditions compared to control conditions. The RDPB is calculated as:

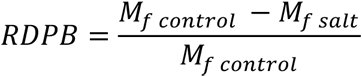

Where M_f_ is plant mass under control and salt treatments as indicated. Lower RDPB values indicate less reduction in biomass under stress conditions and are representative of higher degrees of tolerance. RDPB was converted to percent of plant biomass retained (1-RDPB). Tolerant genotypes were individuals with high amounts of biomass retained, while sensitive individuals retained less biomass in response to treatment.

#### Stress Tolerance Index (STI)

The stress tolerance index (STI, Negrão *et al*., 2017) was calculated for biomass traits (live aboveground biomass, dead aboveground biomass, below ground biomass). The STI was calculated as:

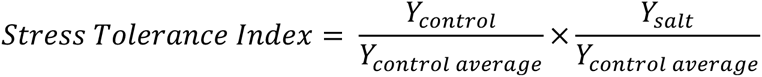

Where Y_control_ and Y_salt_ are measured traits for control and salt treatments for each accession, and Y_control average_ is the trait response under control conditions for the entire population evaluated. A greater STI for an accession indicates higher degrees of salt tolerance. The STI accounts for genotypic response to salinity stress and compares it to a population response to reveal accessions that are performing superior to others. Raw STI values are listed in **Supplementary Table S3**. Raw STI values for live aboveground biomass, dead aboveground biomass, and root biomass were converted to a rank order. STI was rank ordered with 0 indicating missing data, 1 indicating the lowest STI, and 23 indicating the highest possible STI (**Figure 3**).

#### Treatment Effects

Non-metric multidimensional scaling (NMDS) (Julkowska *et al*., 2019), performed in R v. 3.6.0 (R Core Team, 2013), was used to evaluate plant response to salt exposure and to determine groupings among accessions across treatments. The dimcheckMDS function in the geoveg package generated the associated stress value with each reduction in dimension. A lower stress value indicates higher conformity between the true multivariate distance between samples and the distance between samples in reduced dimensions. Two dimensions were deemed appropriate. NMDS was paired with analysis of similarity (ANOSIM), which statistically tests clusters and ordination results from the NMDS. The ANOSIM determines whether the dissimilarity matrix used in the NMDS ordination is significantly different. Using an ANOSIM, we tested treatment effects. Dissimilarities were determined using a Bray-Curtis similarity to test whether accessions were more similar within a treatment compared to among treatments.

#### Landrace and Accession Effects

To determine if plant response to increased salt was a result of genetic mechanisms (accession response or landrace structure), an NMDS was coupled with ANOSIM. The Bray-Curtis dissimilarity coefficients for ST values were used in the NMDS to visualize patterns in the data. Two dimensions were specified. NMDS was paired with ANOSIM to statistically test clusters and ordination results. We tested whether individuals were more similar with an accession compared to among accessions; we tested whether individuals were more similar within a landrace compared to among landraces.

#### Treatment Effects on Growth

One-way analysis of variance (ANOVA) was used to deduce whether there was a statistical difference among accessions for live aboveground biomass STI values, dead aboveground biomass STI values, and root biomass STI values in response to salt exposure. An ANOVA was used to evaluate differences between landraces in response to salt exposure. If significant differences were found, Tukey’s HSD was used to separate accession/landrace means.

Response variables that did not pass a threshold of 0.05 in a Shapiro-Wilk test were transformed and used in the ANOVAs. For the accession ANOVA, STI values for live above ground biomass, dead above ground biomass, and root biomass were square-root transformed. For the landrace ANOVA, STI values for live above ground biomass and dead above ground biomass were log transformed. STI values for root biomass were square-root transformed.

#### Treatment Effects on Sodium and Potassium Accumulation

To determine whether there was significant variation among treatments and accessions with respect to Na^+^ content, K^+^ content, and the potassium to sodium ratio (K^+^/Na^+^), a two-way ANOVA was performed in R v. 3.6.0 (R Core Team, 2013). If a significant difference was found (p<0.05), Tukey’s HSD was performed to determine which treatments and accessions were significantly different from one another.

#### Treatment Effects on Proline Accumulation

To determine whether there was significant variation among treatments and accessions for proline accumulation, a two-way ANOVA was performed on proline values that were log transformed using R v. 3.6.0 (R Core Team, 2013). If a significant difference was found (p<0.05), Tukey’s HSD was performed to determine which treatments and accessions significantly differed from one another.

## RESULTS

### Treatment Effects

Salt exposure reduced live aboveground biomass, root biomass, the shoot-to-root ratio, height, the percent of live leaves, and foliar SPAD across all accessions and landraces, while dead aboveground biomass and mortality increased. *Sorghum* accessions responded differently to NaCl exposure, indicating that variation in salt tolerance exists within our tested population; however, plants were more similar within a treatment rather than across treatments (p<0.001; **Supplementary Fig. S2**).

### Landrace and Accession Effects

Based on accession and landrace ST values calculated for the measured growth parameters (SPAD, percent live leaves, height, mortality, live above ground biomass, dead aboveground biomass, and root biomass), plants were more similar within an accession rather than across accessions (p<0.001) and within a landrace rather than across landraces (p<0.001; **Figure 1**) when exposed to salt, indicating that heritable variation in salt tolerance existed within our tested population.

**Figure 1.**
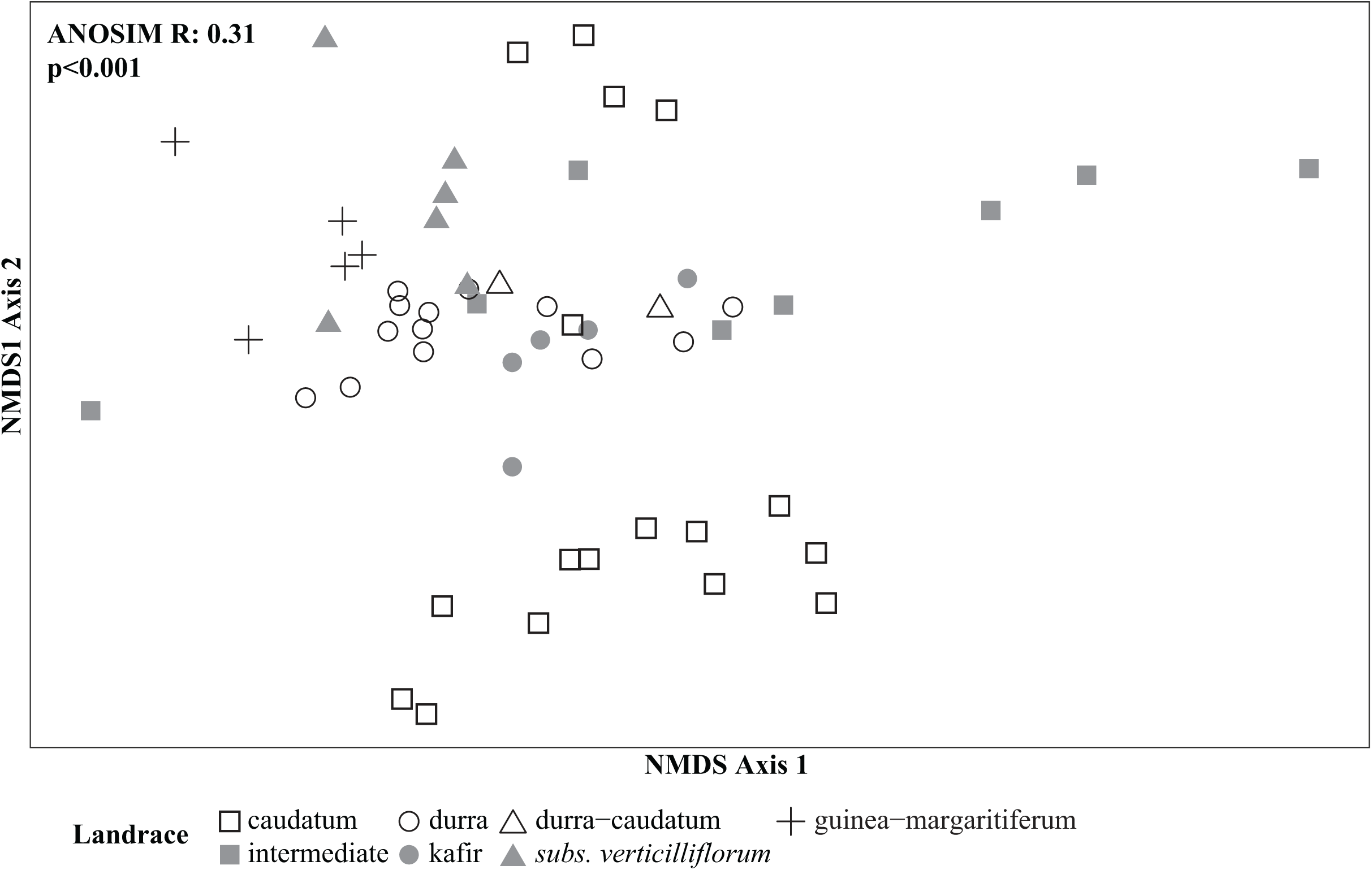
Non-metric multidimensional scaling using Bray-Curtis dissimilarity coefficient to two-dimensionally visualize plant response to treatment. For the NaCl treatment, accessions were ordinated in two-dimensional space. The following measurements were analyzed for dissimilarity among individuals: SPAD, percent of live leaves (live leaf count/total leaf count), height (cm), mortality, live aboveground biomass (dry weight in g), dead aboveground biomass (dry weight in g), and root biomass (dry weight in g). Shapes indicate the landrace grouping for each accession. The analysis of similarity revealed plants were more similar within a landrace than among landraces (R=0.31, p<0.001).

### Relative Decrease in Plant Biomass (RDPB)

Continued growth under stress conditions is an important selective trait for agricultural plant productivity. The percent of biomass retained in response to NaCl ranged from 98% to 3% across accessions (**Figure 2**). Accessions showing sustained growth included V-1 (subs. *verticilliflorum*), Sb-18 (durra), Sb-7 (caudatum), Sb-9 (guinea-margaritiferum), Sb-10 (durra), and Sb-3 (guinea-margaritiferum). These six accessions retained >90% of live aboveground biomass when exposed to NaCl. RDPB values within the NaCl treatment also varied among landraces (p<0.001; **Supplementary Table S1**). High RDPB values, as seen with *S. bicolor* subs. *drummondii* and *S*. *propinquum*, reflect sensitivity to salinity, whereas low RDPB values, as seen with the landrace guinea-margaritiferum, reflect tolerance.

**Figure 2.**
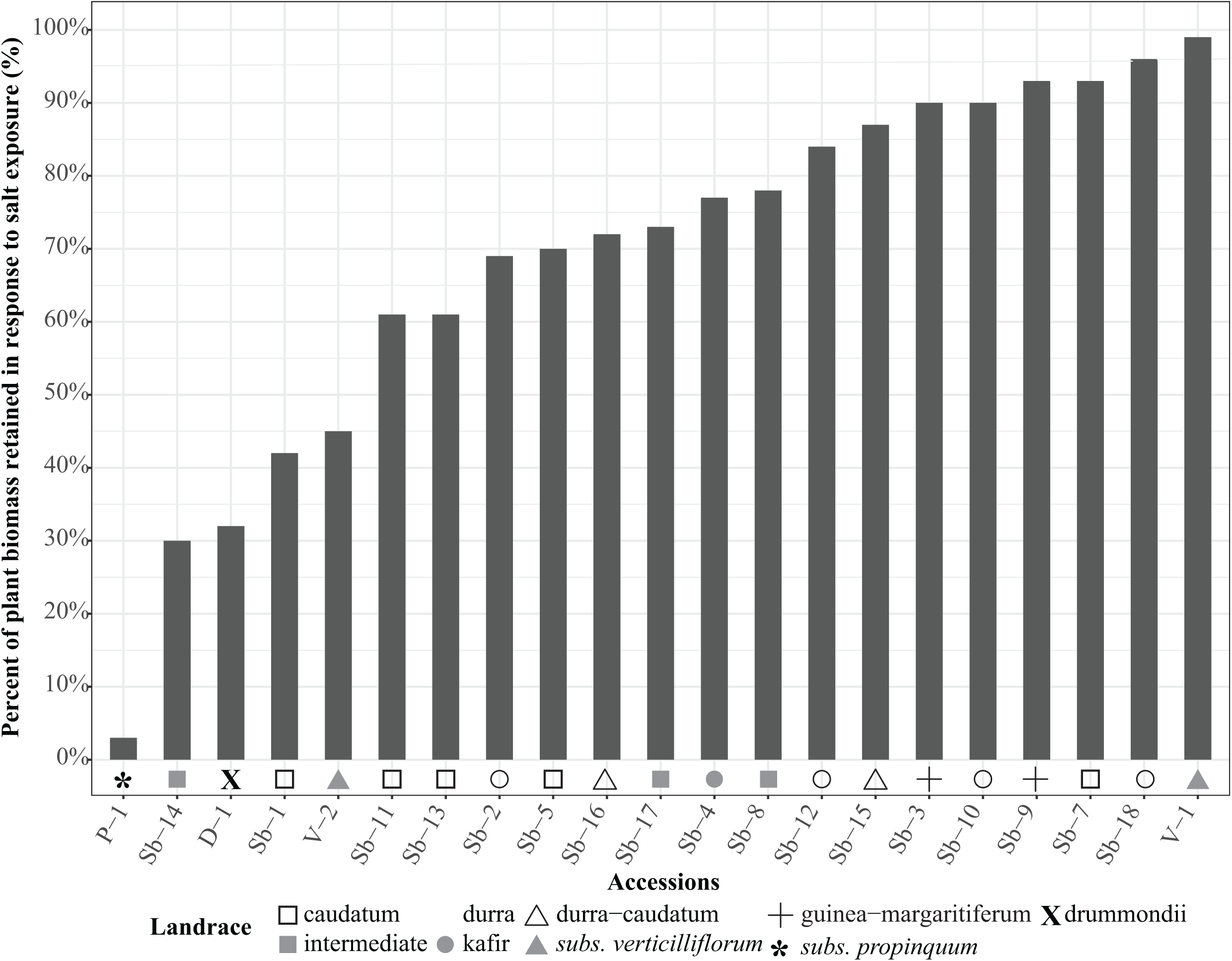
Relative percent of plant biomass retained in response to 75 mM NaCl for each accession. Relative percent of plant biomass retained was calculated by 1-RDPB. Shapes indicate the landrace grouping for each accession. Larger percentages indicate higher amounts of biomass retained in response to NaCl. Lower percentages indicate higher amounts of biomass lost in response to NaCl. RDPB was calculated on mean live above ground biomass in control and treatment conditions.

### Stress Tolerance Index (STI)

The stress tolerance index is a numerical value that describes relative performance of an accession under stress within a population. A larger STI indicates a more tolerant accession compared to others in the population. Raw STI values (**Supplementary Table S2**) for live aboveground biomass, dead aboveground biomass, and root biomass were converted to rank, with larger STI values given a higher rank and lower STI values given a lower rank. STI values for live above ground biomass, dead above ground biomass, and root biomass differed among accessions (p<0.001 for each). STI values ranged from 0.01 to 1.51 for live aboveground biomass, 0.10 to 3.35 for dead aboveground biomass, and 0.05 to 1.97 for belowground biomass. Some accessions ranked high for all three traits while others ranked high for only one or two of the traits. For example, P-1 ranked low for live aboveground biomass (1^st^ out of 21^st^) but ranked 17^th^ out of 21^st^ for root biomass (**Figure 3**), suggesting that, although aboveground biomass was significantly affected in treatment, root biomass was not. The largest overall scores (additive rank score for alive aboveground biomass, dead aboveground biomass, and root biomass) were observed for the accessions Sb-10, V-1, Sb-9, Sb-3, Sb-2, and Sb-12, indicating overall better performance compared to other accessions (**Figure 3**).

**Figure 3.**
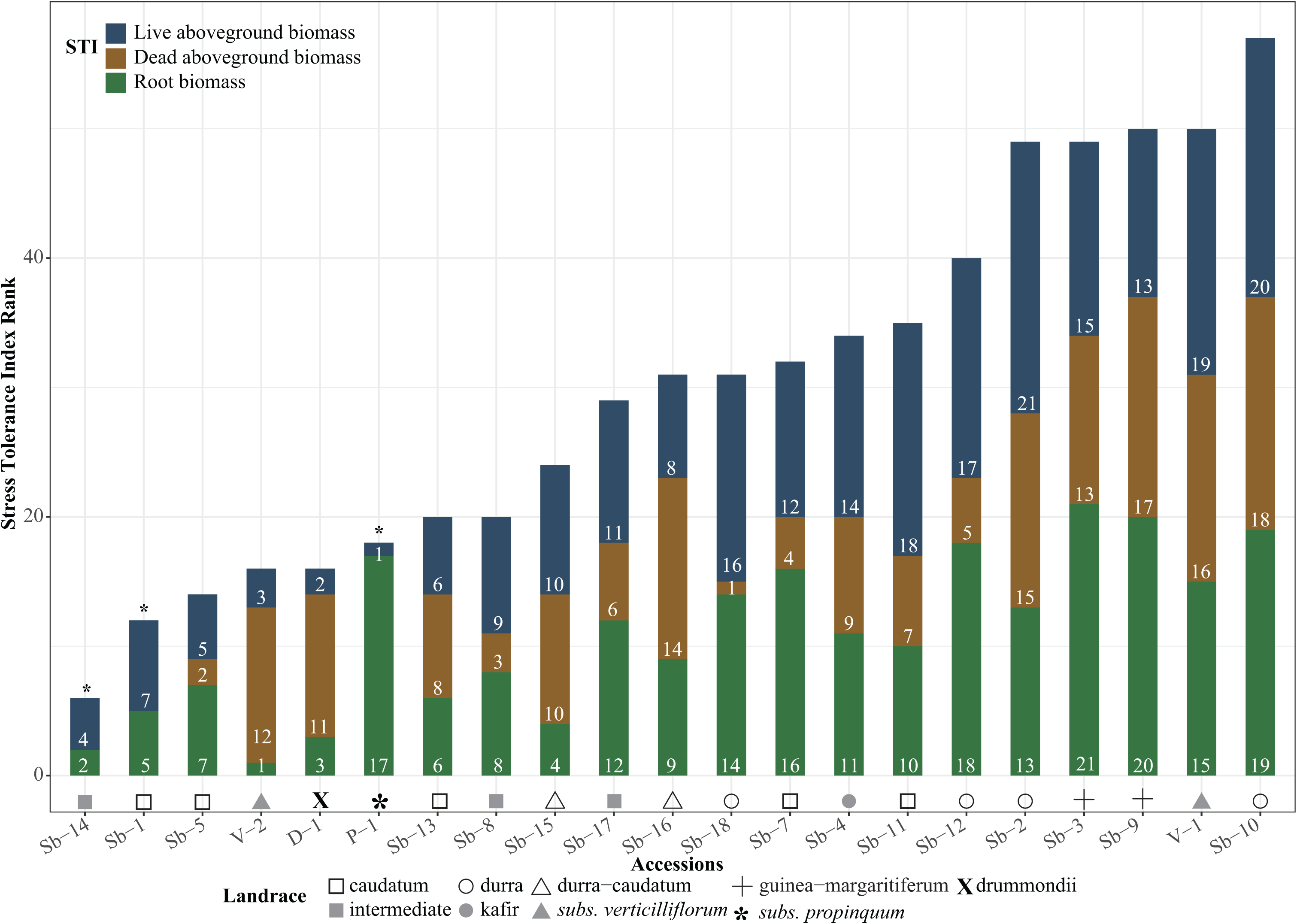
Rank ordered stress tolerance index (STI) scores for live aboveground biomass, dead aboveground biomass, and root biomass, for each accession in response to NaCl. Accessions were arranged with the lowest overall STI rank on the left and the largest overall STI rank on the right. Overall rank was calculated by the sum of live aboveground biomass, dead aboveground biomass, and root biomass rank. Colors indicate portion of overall rank contributed by live aboveground biomass, dead aboveground biomass, and root biomass. Higher values indicate better performers compared to other individuals within the population. Lower values indicate poor performers compared to other individuals within the population. Note: Sb-14, Sb-1, and P-1 are missing STI values for dead aboveground biomass.

When comparing the STI values among landraces, differences were observed for live aboveground biomass, dead aboveground biomass, and root biomass (p<0.001 for each; **Supplementary Table S3**). STI values ranged from 0.01 to 1.28 for live aboveground biomass. *S. propinquum* had the lowest STI for live aboveground biomass with a mean of 0.01 and landrace durra had the highest STI for live aboveground biomass with a mean of 1.28. STI values ranged from 0.32 to 2.08 for dead aboveground biomass with the intermediate landraces displaying the least STI values and the landrace guinea-margaritiferum displaying the highest. STI values ranged from 0.11 to 1.69 for root biomass. The landrace guinea-margaritiferum had the highest STI for root biomass (1.69), while most other landraces averaged about 0.2 to 0.5 (**Supplementary Table S2**).

It is pertinent to point out that the accessions Sb-14, Sb-1, and P-1 are missing data for dead aboveground biomass (noted with an * in **Figure 3**). While this impacts the overall STI rank, as well as individual ranks within each category, this did not hinder our results with respect to tolerance conclusions. Indeed, accessions that ranked as tolerant in the STI analysis overlapped with the accessions that were deemed tolerant in the RDPB analysis. We conclude that we have sufficient data to produce a signal for salt tolerance.

### Sodium and Potassium Accumulation

Significant variation in dead aboveground biomass among accessions (**Supplementary Table S2**) suggests differential Na^+^ accumulation or disruption of K^+^ homeostasis may underlie accession response. A subset of accessions that showed variation in growth under salt treatment were evaluated for ion accumulation. Variation in Na^+^ content was found among treatment and accessions (p<0.001 for each). Foliar Na^+^ under control conditions was low, but varied 35-fold across accessions (**Table 2**). When exposed to NaCl, Sb-3 and Sb-4 accumulated the least amount of Na^+^ while P-1 and V-2 accumulated the most (**Table 2**).

**Table 2.** Summary of Sorghum ion profiles. Sodium (Na^+^), potassium (K^+^), and potassium sodium (K^+^/Na^+^) molar ratios for NaCl treatments for a subset of accessions that showed variability in phenotypic responses. Data shown are means ± (the standard error) of Na^+^ content, K^+^ content, and K^+^/Na^+^ ratio for each accession in the third leaf from the top. Different letters represent significant differences when comparing accessions (p<0.05).

| Accession | Mean Na<sup>+</sup> mg/g |  |  |  | Mean K<sup>+</sup> mg/g |  |  |  | Mean K<sup>+</sup>/Na<sup>+</sup> |  |  |  |
| --- | --- | --- | --- | --- | --- | --- | --- | --- | --- | --- | --- | --- |
|  | Control |  | 75 mM NaCl |  | Control |  | 75 mM NaCl |  | Control |  | 75 mM NaCl |  |
| P-1 | 0.13 | (0.03)<sup>bcdefg</sup> | 2.58 | (0.48)<sup>gh</sup> | 14.32 | (1.54)<sup>bcde</sup> | 3.06 | (1.08)<sup>a</sup> | 81.98 | (25.47)<sup>cdefg</sup> | 0.67 | (0.12)<sup>a</sup> |
| Sb-1 | 0.02 | (0.01)<sup>abc</sup> | 0.59 | (0.16)<sup>efgh</sup> | 17.89 | (1.27)<sup>ef</sup> | 18.18 | (3.03)<sup>ef</sup> | 564.71 | (122.52)<sup>gh</sup> | 23.72 | (6.36)<sup>bcde</sup> |
| Sb-3 | 0.19 | (0.14)<sup>abcde</sup> | 0.15 | (0.07)<sup>abcde</sup> | 9.45 | (0.29)<sup>b</sup> | 14.52 | (0.82)<sup>cde</sup> | 124.51 | (46.12)<sup>cdefg</sup> | 142.67 | (45.08)<sup>defg</sup> |
| Sb-4 | 0.17 | (0.10)<sup>abcdef</sup> | 0.22 | (0.11)<sup>bcdefg</sup> | 13.05 | (0.69)<sup>bcde</sup> | 17.11 | (0.96)<sup>def</sup> | 147.21 | (85.36)<sup>cdefg</sup> | 74.87 | (20.78)<sup>cdefg</sup> |
| Sb-7 | 0.03 | (0.01)<sup>abc</sup> | 0.22 | (0.08)<sup>cdefg</sup> | 22.54 | (1.34)<sup>f</sup> | 22.84 | (0.62)<sup>f</sup> | 806.73 | (240.01)<sup>gh</sup> | 108.48 | (41.20)<sup>cdefg</sup> |
| Sb-8 | 0.04 | (0.01)<sup>abcd</sup> | 0.92 | (0.27)<sup>efgh</sup> | 17.06 | (0.80)<sup>def</sup> | 18.85 | (0.90)<sup>ef</sup> | 246.46 | (41.32)<sup>fgh</sup> | 14.55 | (4.68)<sup>abcde</sup> |
| Sb-9 | 0.03 | (0.01)<sup>abc</sup> | 0.51 | (0.12)<sup>efgh</sup> | 10.93 | (0.85)<sup>bc</sup> | 16.70 | (0.81)<sup>def</sup> | 235.69 | (30.71)<sup>fgh</sup> | 24.12 | (4.75)<sup>bcde</sup> |
| Sb-10 | 0.02 | (0.01)<sup>ab</sup> | 0.60 | (0.23)<sup>defgh</sup> | 10.28 | (0.68)<sup>bc</sup> | 11.43 | (0.92)<sup>bcd</sup> | 703.49 | (466.73)<sup>fgh</sup> | 18.92 | (5.44)<sup>bcd</sup> |
| Sb-15 | 0.09 | (0.07)<sup>abc</sup> | 1.95 | (0.64)<sup>gh</sup> | 18.68 | (0.81)<sup>ef</sup> | 19.34 | (2.23)<sup>ef</sup> | 936.91 | (475.01)<sup>fgh</sup> | 25.15 | (20.61)<sup>abc</sup> |
| Sb-16 | 0.01 | (0.01)<sup>a</sup> | 0.13 | (0.06)<sup>abcde</sup> | 17.86 | (1.22)<sup>ef</sup> | 23.22 | (1.07)<sup>f</sup> | 1569.94 | (674.41)<sup>h</sup> | 351.98 | (179.91)<sup>efgh</sup> |
| Sb-17 | 0.35 | (0.24)<sup>bcdefgh</sup> | 1.79 | (0.74)<sup>fgh</sup> | 14.88 | (0.91)<sup>bcdef</sup> | 18.28 | (0.64)<sup>def</sup> | 94.21 | (65.45)<sup>cdefg</sup> | 9.46 | (4.46)<sup>abc</sup> |
| V-2 | 0.32 | (0.20)<sup>bcdefg</sup> | 3.47 | (1.19)<sup>h</sup> | 11.28 | (1.52)<sup>bcd</sup> | 12.28 | (1.72)<sup>bcde</sup> | 56.12 | (22.40)<sup>cdef</sup> | 2.80 | (0.74)<sup>ab</sup> |
| SEM | 0.45 |  |  |  | 0.08 |  |  |  | 0.45 |  |  |  |
| P<sub>Accession</sub> | p<0.001 |  |  |  | p<0.001 |  |  |  | p<0.001 |  |  |  |
| P<sub>Treatment</sub> | p<0.001 |  |  |  | p<0.050 |  |  |  | p<0.001 |  |  |  |
| P<sub>Interaction</sub> | p<0.001 |  |  |  | p<0.001 |  |  |  | p<0.001 |  |  |  |

As with Na^+^, foliar K^+^ concentrations also varied among accessions and these differed in response to treatments (p<0.001 for all effects). For example, P-1 exhibited relatively low foliar K^+^ under control conditions and then declined more than other accessions under NaCl exposure, whereas K^+^ was unchanged or increased significantly in Sb-3 and Sb-9 under NaCl exposure (**Table 2**).

Maintenance of a high K^+^/Na^+^ ratio is often an indicator of salt tolerant genotypes. In sorghum, we found variation among treatments and accessions for the K^+^/Na^+^ ratio (p<0.001 for all effects). Under control conditions, V-2 had the lowest K^+^/Na^+^ ratio and Sb-16 the greatest. The ratio declined in many accessions under NaCl exposure, most notably in Sb-16 and Sb-15, while the ratio remained relatively high in Sb-3 (**Table 2**).

### Proline Accumulation

In response to salt exposure, proline accumulation in sorghum foliage increased, with the magnitude of increase depending on the accession (p<0.001; **Figure 4**). Proline accumulation ranged from 0.07 to 0.26 gfw^-1^ in the control treatment and 0.07 to 2.63 gfw^-1^ in the salt treatment (**Supplementary Table S4**).

**Figure 4.**
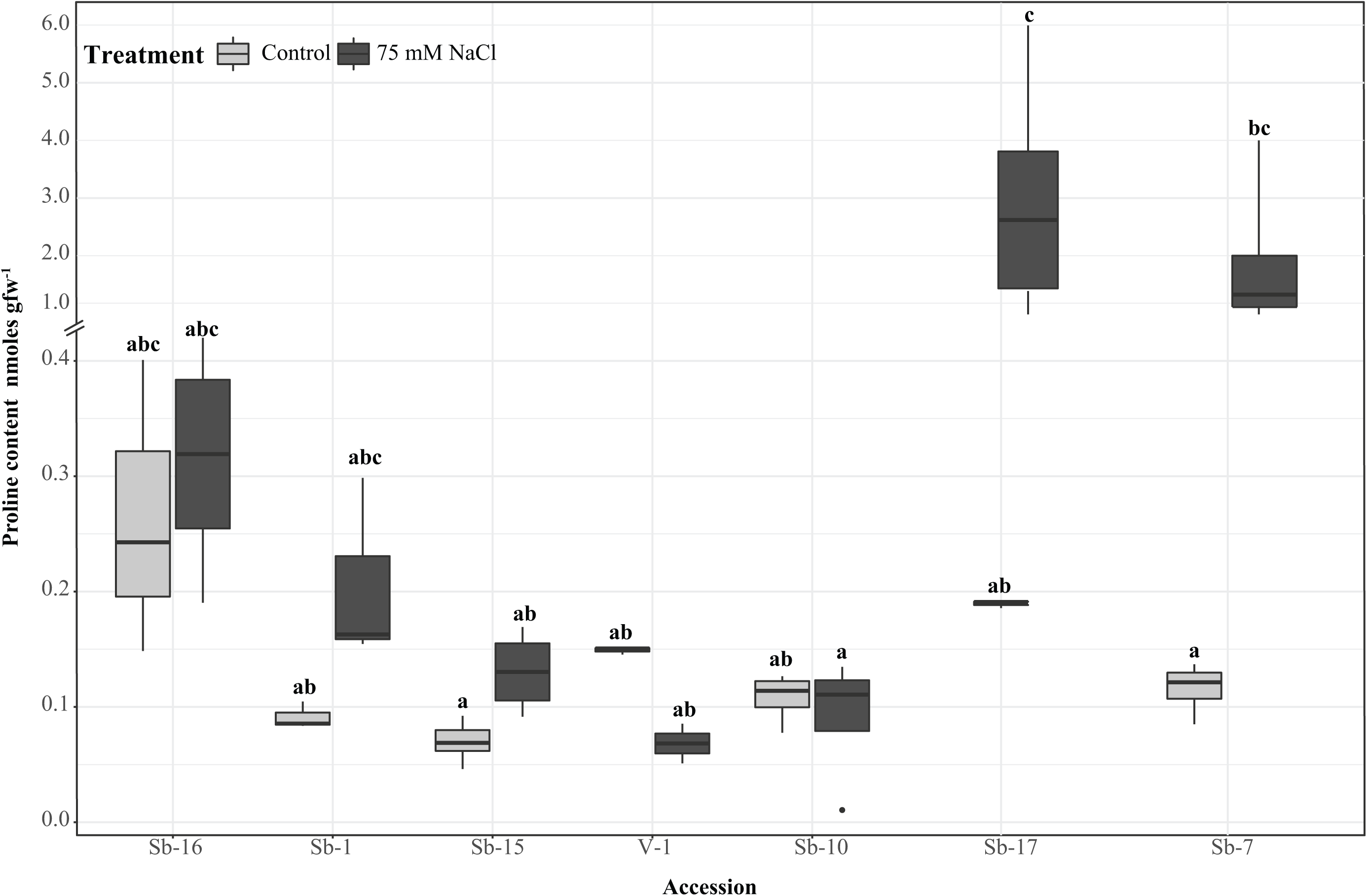
Proline accumulation in a subset of accessions. Some accessions showed no increase in proline accumulation in response to 75 mM NaCl; however, trends for Sb-17 and Sb-7 show that, with increased salt exposure, proline accumulated. Statistical significance was found among accessions and proline accumulation in response to treatment (P_Accession_<0.001, P_Treatment_<0.001, P_Treatment*Accession_ <0.01). Values are the mean of five biological replicates with ± standard error. Different letters represent significant differences. Note: Break in axis to account for scale differences.

## DISCUSSION

### Phenotypic Responses to Salinity Stress

Salinity tolerance is a product of maintenance mechanisms that occur during both the osmotic and ionic phases of salinity stress (Munns & Tester, 2008). During the osmotic phase, continued growth of aboveground biomass indicates the ability to overcome osmotic stress, since sensitivity to water deprivation typically results in decreased growth (Munns & Tester, 2008). In our study, the five accessions with the highest STI values for live aboveground biomass (indicating the ability to obtain sufficient water for sustained growth despite the osmotic impact of salinity exposure) were Sb-2, Sb-10, V-1, Sb-11, and Sb-12 (**Figure 3**).

During the ionic phase, mechanisms of tolerance include compartmentalization of toxic ions into vacuoles and/or extrusion of Na^+^ from cells and the removal of Na^+^ from the xylem stream, which reduces potential exposure in the leaf. Therefore, the accumulation of dead aboveground biomass can be used as proxy for evaluating compartmentalization and extrusion efficiency (Deinlein *et al*., 2014). We find that accessions with high STI values for dead aboveground biomass included both tolerant (Sb-10, Sb-9, and V-1) and sensitive (Sb-16 and V-2) accessions. This, combined with the results for live aboveground biomass, suggests that tolerance in sorghum is correlated to a greater extent with the plant’s ability to overcome the osmotic phase via continued growth rather than exclusion and/or compartmentalization of ions during the ionic phase. This is most evident for accessions such as Sb-10, Sb-9, and V-1. These tolerant accessions accumulated large amounts of both live and dead aboveground biomass (**Figure 3**), reflecting the ability to maintain continued growth under salt exposure.

Plants may exhibit limited root growth as a result of low soil water potential, or conversely, increased growth as a search response for non-saline water. In our study, we found that three of the overall most tolerant accessions (Sb-10, Sb-9, Sb-3) ranked in the top five highest STIs for root biomass (**Figure 3**). This suggests that maintenance of root biomass in response to treatment is associated with salinity tolerance. However, *S. propinquum*, one of the most sensitive accessions, had the largest RDPB and the lowest live aboveground biomass STI, yet had the fifth highest overall root biomass STI (**Figure 2 and Figure 3**, respectively). Given that *S. propinquum* is one of the most sensitive accessions, we conclude that root morphology is not indicative of tolerance. While continued root growth may assist in the search for non-saline water, it does not appear to be a morphological adaptation resulting in tolerance in sorghum.

In our study, we assessed tolerance by relative decrease in plant biomass (RDPB) and the stress tolerance index (STI) (Negrão *et al*., 2017). RDPB is the reduction in growth in response to salt compared to control conditions and is a good measure of the effects of salinity on plant growth within a given accession. The stress tolerance index (STI) is a measurement that accounts for the performance of an individual accession compared to the population under evaluation. We observed that some accessions displayed less than 10% decrease in plant biomass (RDPB) but ranked low in the STI analysis. For example, V-1 and Sb-10 displayed 2% and 10% decreases in plant biomass respectively (or 98% and 90% retained biomass, respectively), in response to treatment and ranked in the top 5 most tolerant accessions in the STI analysis for live above ground biomass, dead above ground biomass and root biomass. However, Sb-18, which lost only 4% of its biomass (retained 96% of its biomass) in response to treatment, ranked 16^th^, 1^st^, and 14^th^ for live above ground biomass, dead above ground biomass, and root biomass respectively. Another example is Sb-7, which lost 7% of its biomass in response to treatment (retained 93%), but ranked 13^th^ overall (12^th^, 14^th^, and 16^th^ for live aboveground biomass, dead aboveground biomass, and root biomass STI, respectively). The discordance between a high rank in the RDPB analysis versus STI analysis suggests that different modes of tolerance may exist in sorghum. Different modes of tolerance may reflect reductions in Na^+^ accumulation achieved by multiple mechanisms, such as reduction in root uptake, reduction in xylem loading, increased extrusion, and increased retrieval from aboveground tissue (Deinlein *et al*., 2014; Wu *et al*., 2019). Each of these mechanisms results in reduced Na^+^ in the cytoplasm. Regardless of the mechanism, reduced Na^+^ typically results in increased tolerance. Therefore, we propose that the RDPB analysis is the better indicator of tolerance because it depicts the outcome of NaCl exposure regardless of the mechanism operating in tolerant genotypes.

### Physiological responses to salinity stress

Historically, proline accumulation under salt and/or osmotic stress has been used as an indicator of tolerance (Iqbal *et al*., 2014). When comparing proline accumulation across accessions, we found that leaf proline increased between the control and NaCl treatment, although this increase was accession dependent (**Figure 4**). V-1 and Sb-10, two of our most tolerant accessions according to RDPB and STI analysis, displayed low amounts of proline in both control and treatment conditions. In contrast, Sb-7 and Sb-17 exhibited large NaCl-induced increases in proline content, but were only moderately salt tolerant. The discordance between proline accumulation and stress tolerance suggests that, in sorghum, proline accumulation may reflect stress injury rather than a mechanism of tolerance. Similarly, other studies have found a lack of correlation between tolerance and proline accumulation. In barley, the QTLs for proline accumulation under stress and for stress tolerance were not linked (Fan *et al*., 2015). In rice, salt-sensitive accessions accumulated higher levels of Na^+^ and proline compared to salt-tolerant accessions (Lutts *et al*., 1999; Vaidyanathan *et al*., 2003; Theerakulpisut *et al*., 2005). Therefore, although proline accumulation does occur in sorghum in response to NaCl, our results suggest that it is not an accurate predictor of protective capacity against stress injury.

Significant variation in sodium and potassium content among accessions suggests that differences in the mechanisms responsible for sodium uptake and distribution and/or regulation of potassium content exist in sorghum (**Table 2**). When comparing the variation in Na^+^ accumulation with tolerance categories, we do not observe patterns suggestive of a unifying mechanism of sorghum response to excess Na^+^. For example, Sb-1 and Sb-10, a sensitive and a tolerant accession, respectfully, did not significantly differ in foliar Na^+^ accumulation. In control conditions both accessions averaged approximately 0.02 mg Na^+^/g, and in treatment conditions both averaged about 0.59 mg Na^+^/g; however, in terms of relative decreases in plant biomass, Sb-10 displayed less than 10% loss in live aboveground biomass while Sb-1 had greater than 50% loss. Although our analysis of foliar Na^+^ by ICP is unable to assess subcellular localization, Sb-10 may have elevated tissue tolerance as a result of better compartmentalization of Na^+^ ions into vacuoles, resulting in less cell death due to ionic imbalance.

Salt sensitivity is often associated with changes in K^+^ uptake resulting from competition between Na^+^ and K^+^ (Deinlein *et al*., 2014). In sorghum, we observed variation in K^+^ among accessions and NaCl treatments. Most variation in K^+^ was observed between accessions and not between treatments. The only accession exhibiting a decline in K^+^ between the control and NaCl treatments was P-1, whereas exposure led to an increase in K^+^ in Sb-3 and Sb-9. Sb-3 and Sb-9 are both from the landrace guinea-margaritiferum and both exhibit low RDPB. In contrast, P-1 had a high RDPB. These patterns suggest that, at least in the sorghum accessions included in this study, the loss of K^+^ homeostasis may not underlie NaCl toxicity, but rather may represent the basis of salt sensitivity in the wild relative, *S. propinquum*.

### Evolution, domestication, and adaptation of salt tolerant sorghum accessions

Where population structure and geographic distribution of sorghum has been studied, landraces show genetic diversity and racial structure with strong geographical patterning (Morris *et al*., 2013; Mace *et al*., 2013). Kafir, which tends to predominate in South Africa, shows the largest genetic variation compared to other landraces, likely due to migration into a contrasting agroclimate (Morris *et al*., 2013). Guinea tends to be widely distributed in western Africa in the tropical savannas. A subgroup of guinea, known as guinea-margaritiferum, is present in the same geographical area, but is understood to have undergone a separate, and more recent, domestication event relative to the other landraces (Morris *et al*., 2013; Mace *et al*., 2013; Mullet *et al*., 2014). Caudatum, primarily found in central-west Africa in tropical savanna climates, displays the least amount of population structure due to exposure to adjacent and varying climates (Morris *et al*., 2013; Mullet *et al*., 2014). Lastly, durra is distributed in warm semiarid deserts in northern Africa and India (Morris *et al*., 2013; Mullet *et al*., 2014). Wild sorghum is known to contain greater genetic diversity compared to landraces, and each landrace was developed through *S. bicolor* outcrossing with wild sorghum in various regions, ultimately resulting in phenotypically diverse plants due to regional adaptation (Kimber, 2000).

We found that salinity tolerance was not solely associated with landrace, suggesting that accessions exposed to high local and regional soil salt contents may have adapted mechanisms to overcome the stresses associated with NaCl exposure. We therefore initially hypothesized that the driving force of variation in salt tolerance may be a result of post-domestication adaptation to saline environments; however, when we evaluate our findings within the phylogenetic framework presented in Mace *et al*. (2013), we observe that the most tolerant *S. bicolor* accessions are those that originated shortly after the domestication event, particularly those accessions within the durra clade (Mace *et al*., 2013, Figure 1, green squares). Further, the two *S. verticilliflorum* accessions included in both this study and the Mace *et al*. (2013) study displayed significantly different responses to salinity. V-1 (PI226096), which had the lowest RDPB and ranked 5^th^ largest for live aboveground biomass STI, dead aboveground biomass STI, and root biomass STI, is positioned in the first post-domestication clade (Mace *et al*., 2013, Figure 1, red triangles); however, V-2 (PI300119), which lost approximately 70% of its biomass in response to treatment and ranked in the 3^rd^ to last position for live aboveground biomass and the last position for root biomass, is placed in the clade prior to the domestication event. This, combined with the observations for the durra accessions, indicates that salinity tolerance was gained during or shortly after sorghum domestication. In contrast, accessions from the landrace caudatum, which displayed a diversity of stress tolerance rankings (**Figure 3**), are not monophyletic, and are found in diverse positions throughout the tree. Interpretation of these results within this phylogenetic context suggests that, during further selection and improvement, salinity tolerance was lost in lineages that were no longer subjected to continued environmental pressure. Lastly, given that *S. bicolor* and especially the landrace durra (Smith *et al*., 2019) is known to be relatively drought tolerant (Mullet *et al*., 2014; Fracasso *et al*., 2016; McCormick *et al*., 2018; Guo *et al*., 2018) and, as with drought stress, salt stress has an initial osmotic component, we propose that salinity tolerance in sorghum originated in combination with, or as a by-product of, drought tolerance during domestication.

## CONCLUSIONS

With more than 500 million people relying on food, fuel, and fiber production from sorghum (Mace *et al*., 2013), the standing genetic diversity of this staple crop should be utilized to maximize production needs, especially in adverse soils. Because of its ability to thrive in environments associated with high degrees of abiotic stressors, it is imperative that the genetic, physiological, and morphological responses to salt exposure in sorghum are understood and utilized to enhance production on saline soils. We identified significant variation in response to salinity exposure among a diverse group of sorghum accessions and we conclude that the variation seen in tolerance is not due to landrace alone, but rather a byproduct of domestication and improvement. Given our results, and in combination with results of Mace *et al*. (2013), we propose that accessions from the landrace durra would serve as valuable resources for genetic improvement of sorghum salinity tolerance in agriculture.

## ACKNOWLEDGEMENTS

The authors wish to thank Dr. Jeffrey Bennetzen for providing *Sorghum propinquum* seed, Dr. Stephen DiFazio for guidance in project design, and Janna Kleinsasser, Natalie Nedley, Rachel Bainbridge, and Margo Folwick for their assistance in data collection. We acknowledge the Pennsylvania State University Analytical Laboratory, State College PA, U.S. National Plant Germplasm System for supplying seed and the West Virginia University Evansdale Greenhouse for supplying space. This work was partially funded by the Eberly College of Arts and Sciences research award (West Virginia University) and the Biology Graduate Student Association graduate student research award (West Virginia University), both awarded to Ashley N. Henderson.

## Abbreviations

RDPB: relative decrease in plant biomass
ST: stress tolerance
STI: stress tolerance index

## SUPPLEMENTARY DATA

**Supplementary Figure S1.** A pilot study showing the effect of increasing concentrations of NaCl on biomass accumulation.

**Supplementary Figure S2.** Non-metric multidimensional scaling using Bray-Curtis dissimilarity coefficient to two-dimensionally visualize plant response to 0 mM and 75 mM NaCl. The analysis of similarity revealed that plants were more similar within a treatment than across treatments (R=0.11; p<0.001). Gray triangles represent individuals within the control treatment and black circles represent individuals within the 75 mM NaCl treatment.

**Supplementary Table S1.** Relative decrease in plant biomass (RDPB) for each landrace. Data shown are means ± the standard error of RDPB values for each landrace. Different letters represent significant differences when comparing landraces (p<0.05).

**Supplementary Table S2.** Accession STI scores and growth variation in response to NaCl. LAGB, live aboveground biomass; DAGB, dead above ground biomass; STI, stress tolerance index; RB, root biomass. Data shown are means ± the standard error. Different letters represent significant differences when comparing accessions (p<0.05).

**Supplementary Table S3.** Landrace STI scores and growth variation in response to NaCl. LAGB, live aboveground biomass; DAGB, dead above ground biomass; STI, stress tolerance index; RB, root biomass. Data shown are means ± the standard error. Different letters represent significant differences when comparing landraces (p<0.05).

**Supplementary Table S4.** Mean proline content for control and NaCl conditions. Data shown are means ± the standard error of proline (gfw-1) for a subset of accessions. Different letters represent significant differences when comparing accessions and treatment (p<0.05).

