## Supplemental Figure S2 for "Phenotypic and physiological responses to salt exposure in *Sorghum* reveal diversity among domesticated landraces"

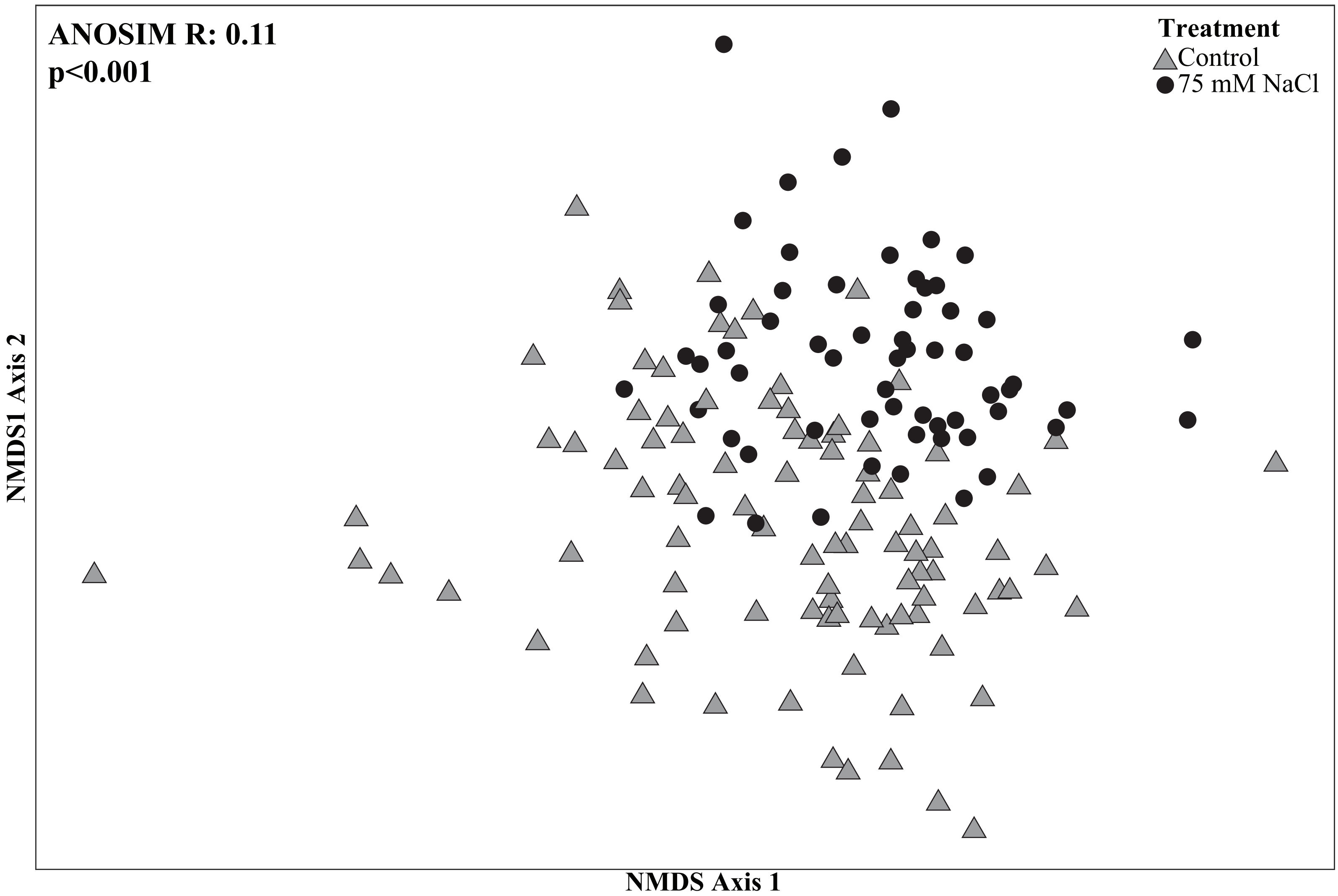


**Supplementary Figure S2.** Non-metric multidimensional scaling using Bray-Curtis dissimilarity coefficient to two-dimensionally visualize plant response to 0 mM and 75 mM NaCl. The analysis of similarity revealed that plants were more similar within a treatment than across treatments ($R=0.11$; $p<0.001$). Gray triangles represent individuals within the control treatment and black circles represent individuals within the 75 mM NaCl treatment.
