## Supplemental Figure S1 for "Phenotypic and physiological responses to salt exposure in *Sorghum* reveal diversity among domesticated landraces"

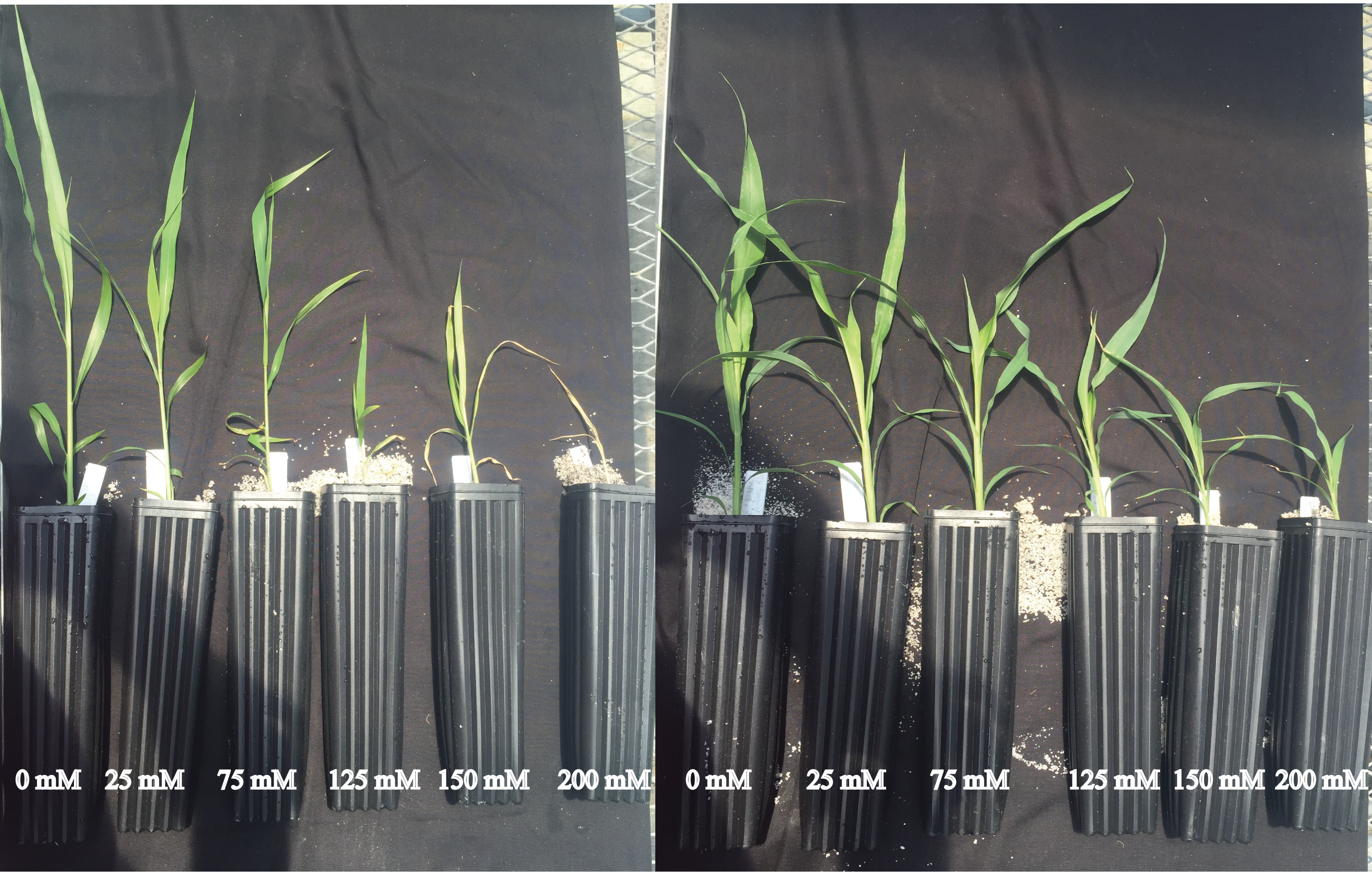


**Supplementary Figure S1.** A pilot study showing the effect of increasing concentrations of NaCl on biomass accumulation.
